# NET-prism enables RNA polymerase-dedicated transcriptional interrogation at nucleotide resolution

**DOI:** 10.1101/246827

**Authors:** Constantine Mylonas, Peter Tessarz

## Abstract

The advent of quantitative approaches that enable interrogation of transcription at single nucleotide resolution has allowed a novel understanding of transcriptional regulation previously undefined. However, little is known, at such high resolution, how transcription factors directly influence RNA Pol II pausing and directionality. To map the impact of transcription/elongation factors on transcription dynamics genome-wide at base pair resolution, we developed an adapted NET-seq protocol called NET-prism (<u>N</u>ative <u>E</u>longating <u>T</u>ranscription by <u>P</u>olymerase-<u>R</u>egulated Immunoprecipitant<u>s</u> in the <u>M</u>ammalian genome). Application of NET-prism on elongation factors (Spt6, Ssrp1), splicing factors (Sf1), and components of the pre-initiation complex (PIC) (TFIID, and Mediator) reveals their inherent command on transcription dynamics, with regards to directionality and pausing over promoters, splice sites, and enhancers/super-enhancers. NET-prism will be broadly applicable as it exposes transcription factor/Pol II dependent topographic specificity and thus, a new degree of regulatory complexity during gene expression.

## INTRODUCTION

Transcription is a highly dynamic process that comprises three different stages. Initiation involves RNA Polymerase II recruitment to the promoter followed by release of RNA Pol II towards progressive elongation. Transcriptional termination is promoted when RNA transcripts are processed and RNA Pol II is released from the chromatin template (Harlen and Churchman, 2017; Jonkers and Lis, 2015). This dynamic shift from one stage to another is facilitated by a compendium of regulatory processes involving phosphorylation of the Pol II C-terminal domain (CTD) and recruitment of factors that facilitate and regulate RNA Pol II activity (Bentley, 2014; Jeronimo et al., 2015; Voss and Hager, 2013).

Approaches that precisely map the position of RNA Pol II at a high resolution have provided a deeper insight into transcriptional regulatory mechanisms (Churchman and Weissman, 2011; Fischl et al., 2017; Kwak et al., 2013; Mayer et al., 2015; Min et al., 2011; Scruggs et al., 2015). For example, the development of the human NET-seq protocol quantitatively purifies Pol II in the presence of a strong Pol II inhibitor hence omitting the utilisation of an antibody (Mayer et al., 2015). Although, this particular approach successfully maps the 3’end of nascent RNA to reveal the strand-specific position of Pol II with single nucleotide resolution, it does not distinguish between different Pol II variants or specific protein-dependent interactions. A similar protocol, the mammalian NET-seq protocol (mNET-seq) uses an immunoprecipitation step to capture the nascent RNA produced by different C-terminal domain (CTD) phosphorylated forms of Pol II (Nojima et al., 2015). Immunoprecipitation has a potential second benefit as it would allow in principle the interrogation of transcription factor – RNA Pol II interaction genome-wide quantitatively, at nucleotide resolution and strand-specificity, none of which is possible using conventional ChIP-seq. In *S. cerevisiae*, such an approach – coined TEF-seq - was developed to interrogate Paf1 – RNA Pol II interaction (Fischl et al., 2017) and allowed new insight into Paf1 requirements for gene expression.

Here, we sought to develop a mammalian counterpart to TEF-seq, which includes an immunoprecipitation step of RNA Pol II associated factors, while being efficient enough to capture sufficient amounts of nascent RNA for processing the latter as part of NET-seq type libraries. A detailed protocol is outlined in Figure 1A and also in a step-by-step form as part of the Supplementary Information (“NET-prism protocol”).

**Figure 1:**
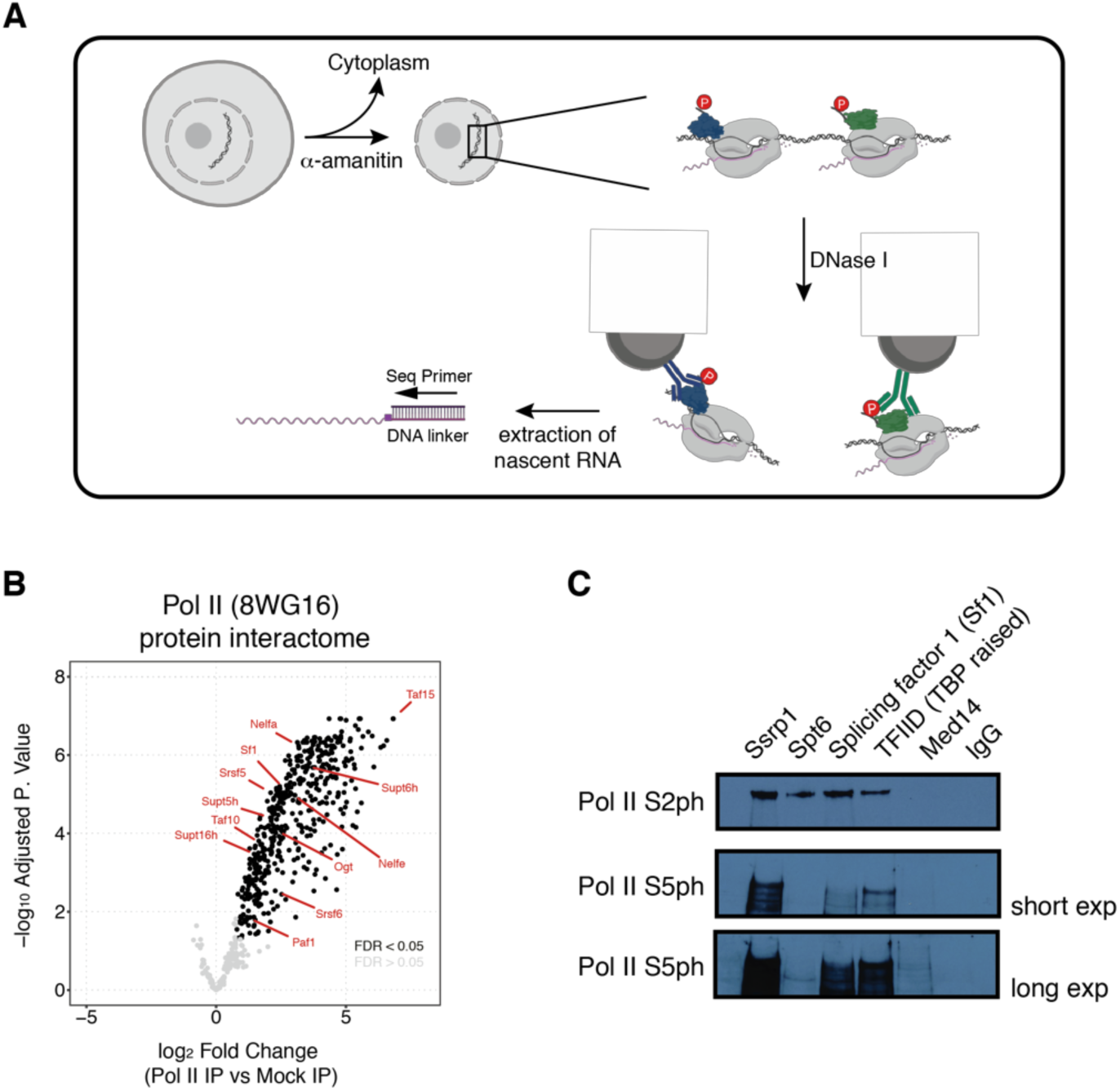
NET-prism as a tool to interrogate active RNA Pol II – interaction with associated proteins. (A) Schematic representation of the approach. For detailed experimental conditions, please refer to the Supplementary Information (“NET-prism protocol”). Upon extraction of nascent RNA, libraries are made using the human NET-seq protocol (Mayer et al., 2015). (B) Volcano plot of Pol II IP vs Mock (IgG) IP depicting a whole RNA Pol II – protein interactome as assessed by Mass spectrometry. Significant values (FDR < 0.05) are coloured in black. (C) Independent confirmation of identified RNA Pol II interactors using IP conditions as used in NET-prism followed by western blotting.

## RESULTS

### Extraction conditions of NET-prism allow for co-purification of RNA Pol II with known associated complex members

Similarly to the original yeast NET-seq and TEF-seq protocols (Churchman and Weissman, 2011; Fischl et al., 2017), we relied on a strong inhibitor for Pol II (α-amanitin) to prevent run-on of the polymerase during all lysis steps and on DNase I to solubilize chromatin. We optimized conditions for DNase I treatment in the absence and presence of urea, which proved to be necessary for efficient solubilization. We found that 100U DNase I and 50mM urea were sufficient to release a large fraction of engaged RNA Pol II from chromatin (Supplementary Figure 1). We then wanted to investigate if we were able to IP known and new co-factors of RNA Pol II under these experimental conditions and examined the total Pol II protein interactome by Mass spectrometry using the same extraction conditions as NET-prism to identify such factors in a native chromatin state.

We identified both, positive (Supt5, Supt6, FACT, Paf1) and negative (NELF) elongation factors as well as splicing (Srsf5, Srsf6, Sf1) and TFIID (Taf10, Taf15) components as significantly enriched with Pol II under NET-prism conditions (Figure 1B and Supplementary Table 1), equipping researchers with a list to guide any follow-up experimentation. We were also able to confirm some of these interactions using immunoprecipitation followed by western blotting (Figure 1C).

### NET-prism captures unique transcriptional footprints of RNA Pol II-associated factors

We picked one transcription factor, TFIID (antibody raised against TBP) and two elongation factors, Spt6 and Sssrp1 (subunit of the FACT heterodimer) to validate the NET-prism approach and interrogate the impact of these factors on RNA Pol II activity. We also performed an IP for Mediator (Med14), serving as a negative control, since it did not display a significant association with Pol II under the conditions used to solubilise chromatin (Figure 1C). The data were highly reproducible among biological replicates (Supplementary Figure 2A) and exhibited diverse correlations with total RNA Pol II over promoter regions (Supplementary Figure 2B – NET-seq/prism), indicating that different TFs establish unique RNA Pol II footprints. Indeed, aligned and averaged NET-prism profiles over the TSS demonstrate additional regulatory complexity during transcriptional initiation and elongation, suggesting that TF binding specificity directly affects RNA Pol II initiation and elongation dynamics. IPs for elongation factors Spt6 and Ssrp1 show strong and broad enrichment of the Pol II complex. These data are in agreement with ChIP-seq densities for both elongation factors (Wang et al., 2017). On the other hand, TFIID-bound RNA Pol II displays a sharp signal centred around the TSS, whereas an IP for Mediator (Med14) yields no nascent RNA transcripts as there is minimal interaction between Mediator and RNA Pol II under the conditions used (Figure 2A). Similar RNA Pol II patterns were also confirmed at a single gene level (Figure 2B). To test more systematically how different transcription factors might influence RNA Pol II initiation and elongation, we sought to determine whether different NET-prism libraries provide improved resolution of RNA Pol II distribution patterns. We calculated the travelling ratio (TR) in the sense direction which is defined as the density of RNA Pol II over the promoter (−30 to +250 bp around the TSS) versus the gene body area (+300 bp downstream of the TSS to −200 bp upstream of the TES). All NET-prism libraries exhibited different TRs indicating different pause-release dynamics of Pol II when bound by different TFs (Figure 2C) These data suggest that NET-prism is indeed able to resolve mechanistic and dynamic interplays between transcription factors and active RNA Pol II. Nascent RNA transcripts upstream of the TSS (sense direction) are too short to produce mappable sequencing reads as the minimum read length is ∼18 nt for unique alignment to the mammalian genome. Therefore, in order to characterise Pol II distribution over those sites we specifically examined anti-sense transcription and coined the “Reverse travelling ratio” which is defined as the density of initiating divergent Pol II versus the density of elongating divergent Pol II (Supplementary Figure 2C). We observed different reverse travelling ratios for all TFs that are bound to RNA Pol II further reinforcing the unique propensity of each TF in controlling anti-sense pause release.

**Figure 2:**
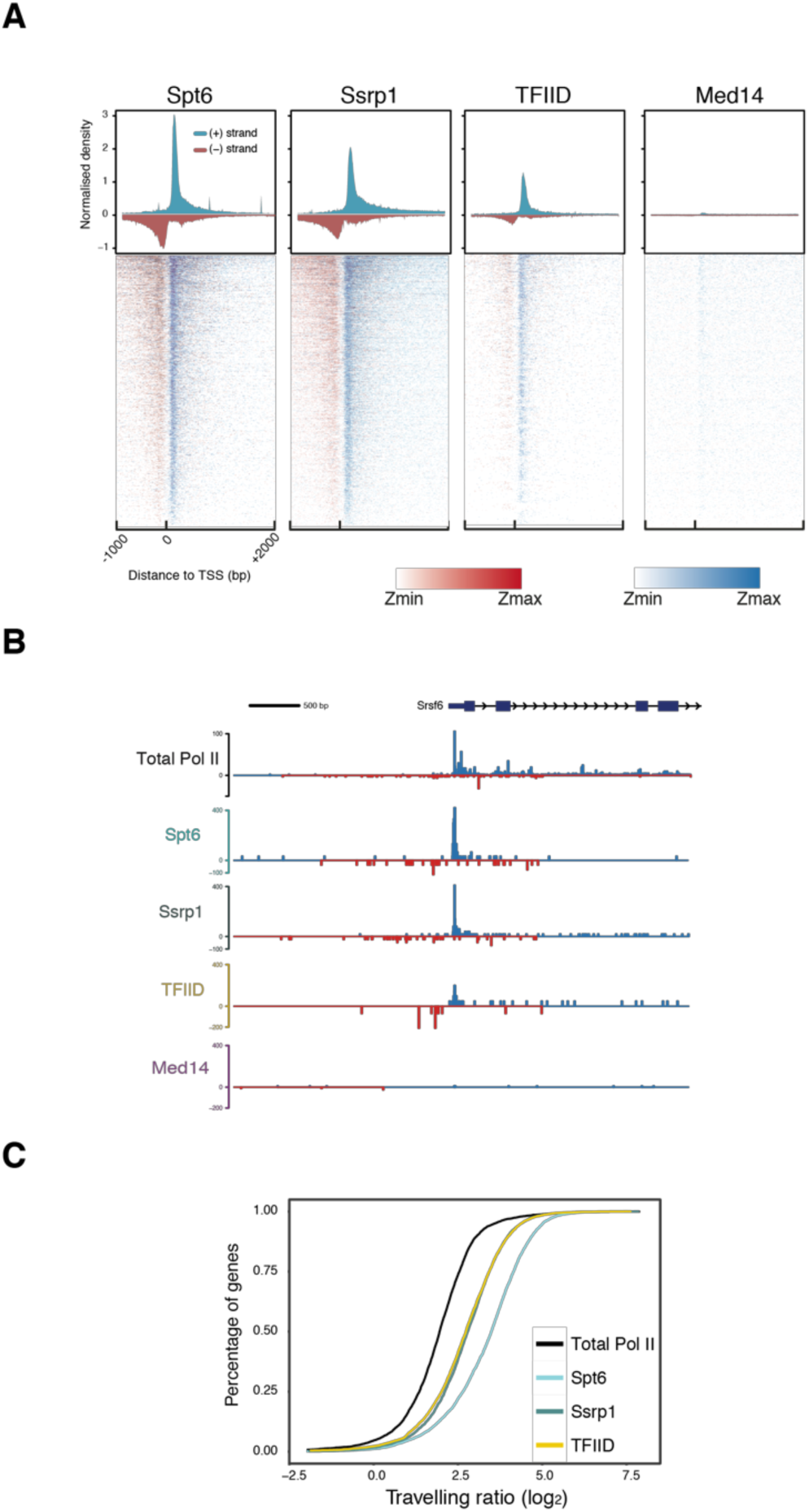
NET-prism application on polymerase-bound transcription factors. (A) Metaplot profiles and heatmaps over protein-coding genes (n =4,314) for polymerase associated elongation (Spt6, Ssrp1) and initiation (TFIID, Med14) factors. A 10-bp smoothing window has been applied. Blue = Sense transcription, Red = Anti-sense transcription. (B) RNA Pol II interrogation of all NET-seq/prism libraries over a single gene (*Srsf6*). (C) Cumulative distribution of Pol II travelling ratio as assessed by NET-prism.

Interestingly, as all three factors examined here (Spt6, SSRP1 and as TFIID antibody was raised against TBP) also interact with RNA Pol I and III, we asked if nascent transcript would also stem from gene products of these two polymerases. Indeed, all three IPs against these proteins also pull-down nascent transcripts generated by RNA Pol I and Pol III (Supplementary Figure 2D), suggesting that NET-prism might be an ideal tool for the investigation of all three RNA polymerases.

### Sequential NET-prism confirms that nascent RNA stems from direct interaction between active RNA Pol II and Ssrp1

Using western blotting, we showed that the conditions of NET-prism allow co-purification of RNA Pol II in the IPs against transcription factors (Figure 1C). However, at least one of them, Ssrp1 has been previously reported to bind RNA (Baltz et al., 2012; Beckmann et al., 2015; He et al., 2016). Therefore, we decided to test more rigorously if the recovered nascent RNA is specifically associated with RNA Pol II and does not stem from direct binding of Ssrp1 to nascent RNA. In order to address this question, we performed a sequential IP as part of NET-prism as outlined in Figure 3A.

**Figure 3:**
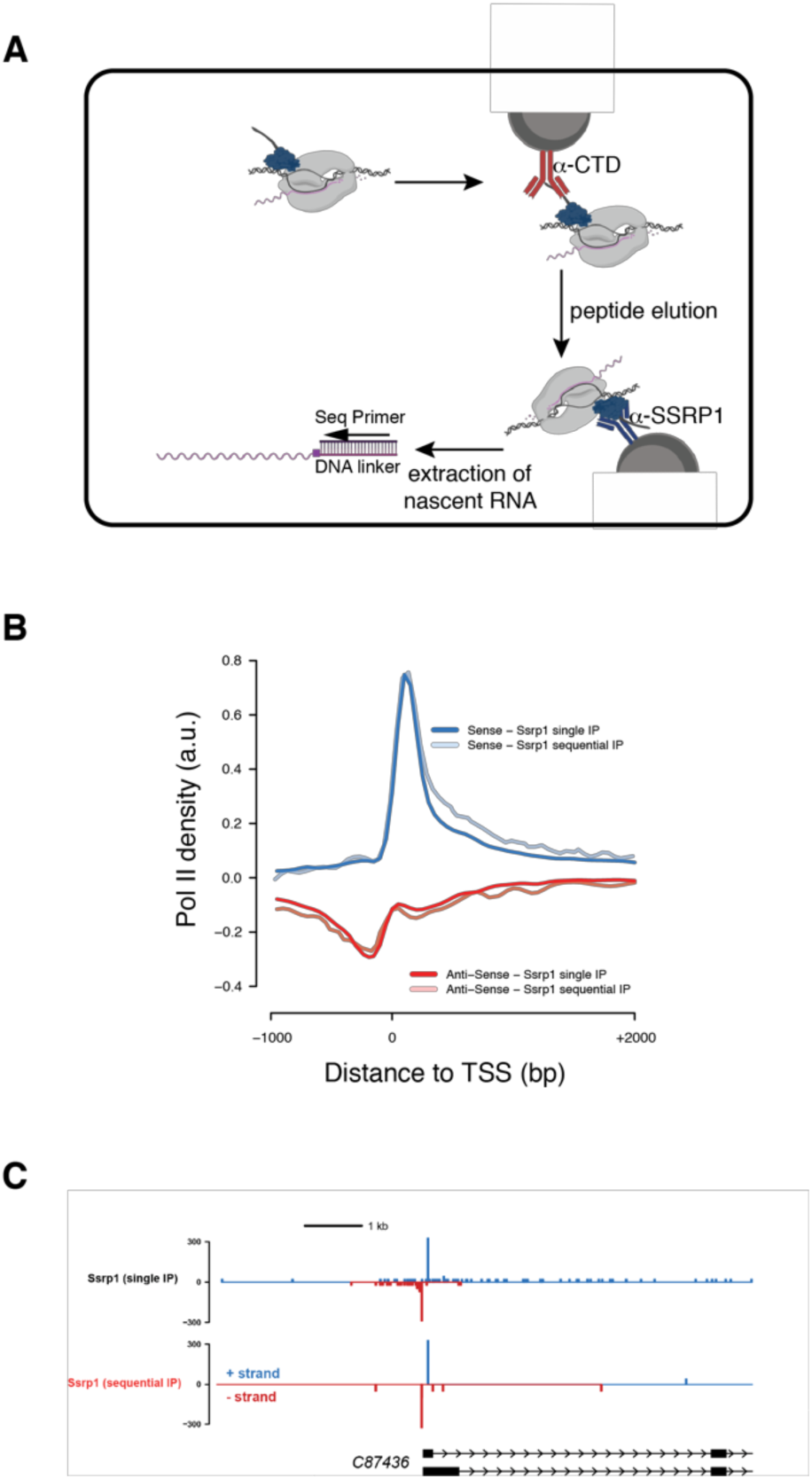
Sequential NET-prism. (A) Schematic illustration of the experiment. Peptide elution was performed using the same CTD peptide used for antibody production and was used in excess. (B) Metaplot profile comparing single and sequential IP. (C) Single gene snapshot comparing single and sequential IP.

Initially, RNA Pol II was immunoprecipitated using an anti-CTD antibody, followed by competitive elution of RNA Pol II by an excess of CTD peptide (the exact peptide used to generate the α-CTD antibody). The eluent subsequently served as input for the second round of IP using an anti-SSRP1 antibody to capture exclusively SSRP1-bound Pol II complexes. The isolated nascent RNA was subsequently used for library generation. Importantly, comparing single and sequential IP by metagene profiling (Figure 3B) and single gene interrogation (Figure 3C) revealed high similarity, strongly suggesting that indeed, NET-prism captures only nascent RNA bound by RNA polymerases and not by TFs.

### NET-prism reveals high resolution Pol II pausing at intron-exon boundaries

Transcriptional elongation rates can affect splicing outcomes suggesting that transcription and splicing are tightly coupled (Dujardin et al., 2014; Fong et al., 2014). Data generated by human NET-seq, mNET-seq, and PRO-seq are consistent with this kinetic model of splicing regulation (Kwak et al., 2013; Mayer et al., 2015; Nojima et al., 2015). While mNET-seq already implicated different RNA Pol II variants to play distinct roles during splicing dynamics (Nojima et al., 2015), it is not known, whether transcription (elongation) factors facilitate RNA Pol II pausing at splice sites. We also reasoned that NET-prism might be an ideal tool to dissect splicing factor – RNA Pol II interaction at splice sites. Therefore, we performed an additional NET-prism library for Splicing factor 1 (Sf1) and included this in our splicing dynamics analysis. As splicing intermediates are known NET-seq contaminants due to the presence of 3’-OH groups in these RNAs (Mayer et al., 2015), we removed them to avoid bias. For the splicing dynamics analysis, we assessed total RNA Pol II (NET-seq (Mylonas et al., 2018)) and NET-prism data for Ssrp1, Spt6, TFIID and Sf1 over intron-exon boundaries. Total RNA Pol II in mouse ES cells showed increased pausing at exon boundaries similarly to human cells (Mayer et al., 2015) (Figure 4A – Total Pol II). Exploration of NET-prism datasets confirmed that only Sf1 exhibited similar pausing at exon boundaries (Figure 3A & Supplementary Figure 4A). In addition, components of the PIC did not associate with Pol II pausing over spliced sites (Supplementary Figure 4B). Interesting to note is also the fact that NET-prism libraries displayed higher Pol II density over exons as opposed to introns suggesting that transcriptional elongation is slower at exons in mouse ES cells (Figure 4B,C). In addition, Sf1-PolII interaction clearly marks exons, indicating specificity of the approach. Taken together, these results augment the kinetic model of transcription and splicing coupling. Our data in combination with previously published results therefore suggest that transcriptional splicing mechanics is facilitated by Pol II variants and elongation factors differently and NET-prism might represent one ideal tool to address this at high resolution.

**Figure 4:**
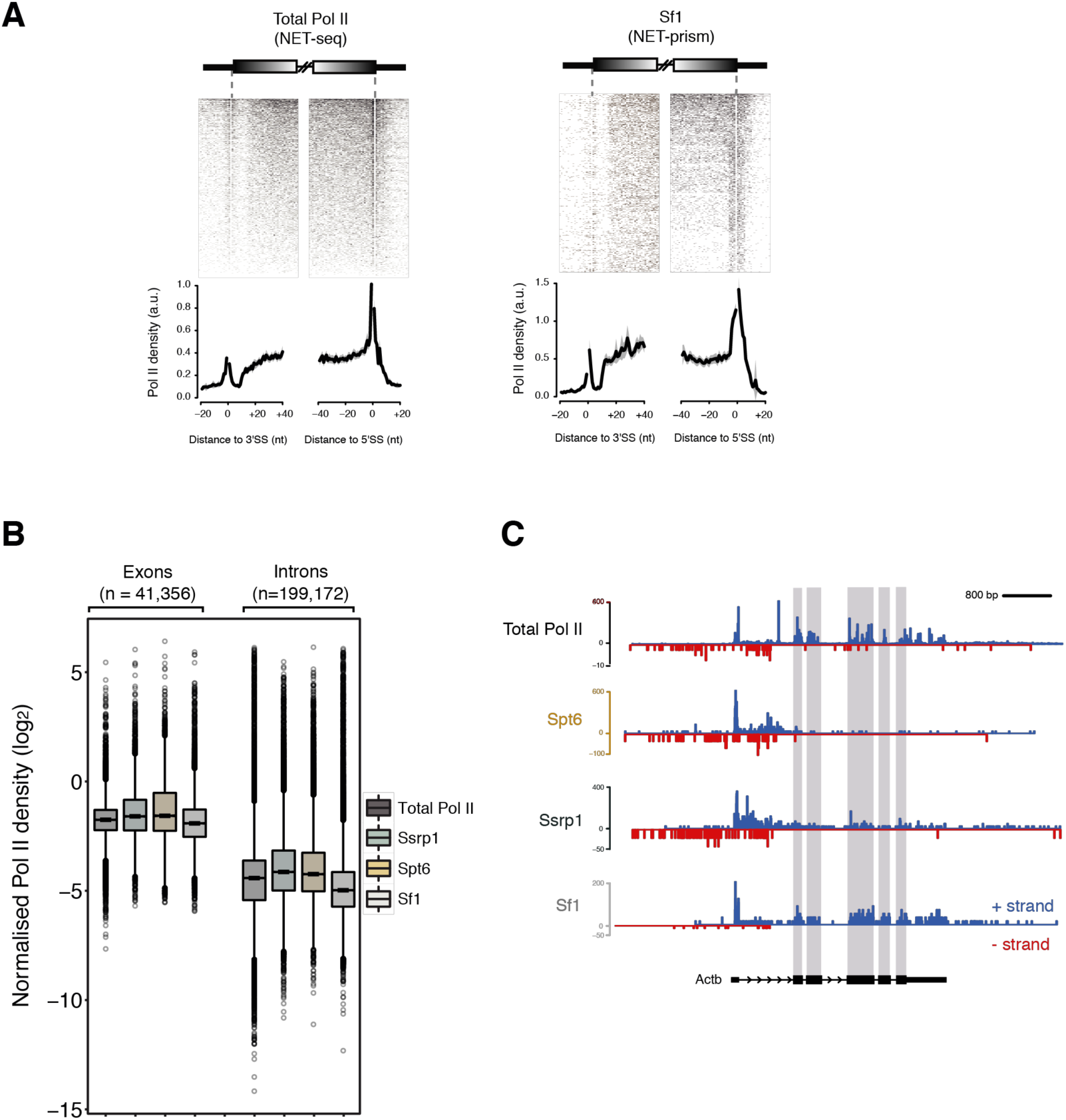
Association of different proteins with transcriptional splicing as assessed by NET-prism. (A) Heatmaps and metaplots assessing polymerase pausing for total Pol II and Splicing factor 1 (Sf1) over exon boundaries (n = 5,550). Solid lines indicate the mean values, whereas the shading represents the 95% confidence interval. (B) Boxplots measuring Pol II coverage over exons (n = 41,356) and introns (n = 199,172) for each NET-seq/prism library. First and last exons are removed from the analysis. (C) RNA Pol II interrogation of all NET-seq/prism libraries over a single gene (*Actb*). Exons are highlighted in purple.

### NET-prism reveals diverse transcriptional dynamics at enhancers

Enhancers and super-enhancers have been shown to play a prominent role in the control of gene expression programs essential for cell identity across many mammalian cell types (Adam and Fuchs, 2016; Heinz et al., 2015; Whyte et al., 2013). Production of enhancer RNAs (eRNAs) is bidirectional and is governed by distinctive patterns of chromatin accessibility (Core et al., 2014), but it is not well characterised whether the same transcriptional rules apply over enhancers as in promoters, in terms of initiation and elongation. We therefore extended our analysis to identify high resolution Pol II footprints at distal and super-enhancers using NET-prism. Highest correlations were identified among Total Pol II and Ssrp1 both for distal and super-enhancers (Figure 5A). Total Pol II and TFs exhibited significantly higher ChIP-seq density over super-enhancers as opposed to distal enhancers. Concomitantly, increased transcriptional activity was confirmed over super-enhancers via NET-prism suggesting TF density being proportional to the degree of Pol II recruitment (Figure 5B). Strikingly, both metaplot profiling (Supplementary Figure 5) and single enhancer (Figure 5C) interrogation of NET-prism transcriptional activity exposed distinctive topographic footprints; Ssrp1 displayed patterns similar to transcriptional initiation whereas Spt6 imitated a trail reminiscent of transcriptional elongation. Moreover, transcriptional activity prompted by TFIID also supports, to some degree, a notion of transcriptional initiation over enhancers (Figure 5C).

**Figure 5:**
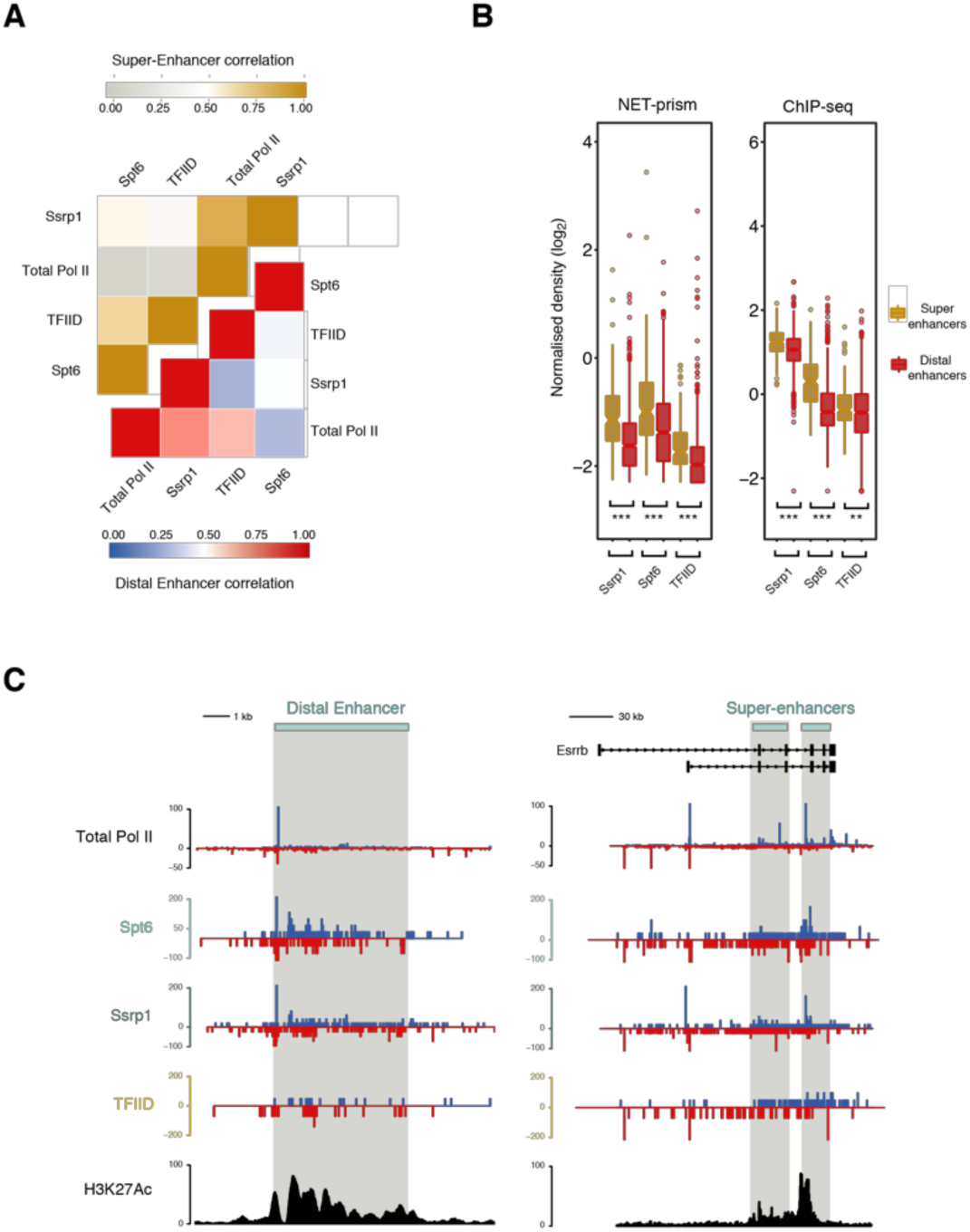
Distinctive patterns of transcriptional regulation over enhancers and super-enhancers. (A) Pearson’s correlation heatmap among NET-seq/prism libraries over distal enhancers (blue – red) and super-enhancers (grey – gold). (B) Boxplots measuring either transcription factor (ChIP-seq) or Pol II (NET-prism) density over distal enhancers (red) and super-enhancers (gold). Significance was tested via the Wilcoxon rank test (** p< 1.0e^-10^, *** p< 2.2e^-16^). (C) Pol II distribution over a distal (chr1: 86,484,171 – 86,495,700) or super-enhancer as assessed by NET-seq/prism. H3K27Ac density is depicted in black colour. Blue and red depict RNA Pol II pausing in the positive and negative strand, respectively.

## DISCUSSION

Here, we have developed a new approach to accurately assess transcriptional topography at a high resolution. In summary, NET-prism allows the direct strand-specific investigation of the transcriptional landscape at single nucleotide resolution of any protein of interest in complex with RNA Pol II. Its robustness enables a deeper insight into the interplay of transcriptional mechanisms conferred by different Pol II variants and proteins that are bound to Pol II. The comprehensive Pol II - protein interactome that we provide here (Supplementary Table 1) facilitates the choice of the protein of interest when applying NET-prism. In addition, given the right RNA polymerase inhibitors and antibodies, NET-prism can be extended to specifically interrogate nascent transcription governed by either RNA Pol I or Pol III.

We hypothesize that NET-prism will be an ideal tool to investigate transcription/elongation factor interactions with actively travelling RNA polymerase at single nucleotide resolution and with strand specificity. An analogous approach has been previously developed in yeast (Fischl et al., 2017), where a variant of the yeast NET-seq protocol (Churchman and Weissman, 2011), called TEF-seq, reveals distinctive patterns of Pol II when bound by diverse elongation factors (Paf1, Spt6, Spt16). Similarly to this approach, NET-prism exposes diverse Pol II signals for every immunoprecipitated TF implying the different dynamics conferred by TF-associated RNA Pol II.

Moreover, our study yields a global picture of how transcriptional elongation is affected at splicing sites and NET-prism might shed light on an unresolved dogma encompassing splicing catalysis. The idea of transcriptional elongation influencing alternative splicing arises from two unique models; the recruitment model (differential recruitment of splicing factors) and the kinetic model (Pol II pausing determines the timing in which splicing sites are presented) (Bentley, 2014; Dujardin et al., 2014). Similarly to other high resolution approaches (Kwak et al., 2013; Mayer et al., 2015; Nojima et al., 2015), we show that splicing is associated with Pol II exon density and strong pauses at both the 3’ and 5’SS, consistent with the kinetic model.

It is important to recognize though that NET-prism – similarly to ChIP-seq – greatly relies on the quality of the antibody used. Antibody cross-reactivity might result in unspecific binding and thus, generation of artefactual RNA Pol II footprints. Therefore, the choice of a highly specific antibody for the protein of interest is important to achieve unique RNA Pol II footprints.

Similarly to the human NET-seq (Mayer et al., 2015), we expect the adaptation of NET-prism to be equally straightforward in any higher eukaryotic cell type. The use of an IP step in NET-prism makes it practical for studying a range of different Pol II - associated factors in order to improve our understanding of transcriptional elongation and its connection to transcript fate. The combination of NET-prism with a high resolution ChIP-seq technique, such as ChIP-nexus (He et al., 2015), could illuminate how exactly *in vivo* binding of transcription or splicing factors correlates with transcriptional activity over different cell states and conditions. Therefore, NET-prism will become a valuable tool for unravelling transcriptional and regulatory complexity.

## MATERIAL AND METHODS

### Cell culture

The E14 cell line (mESCs) was cultured at 37 °C, 7.5% CO_2,_ on 0.1% gelatin coated plates, in DMEM + GlutaMax™ (Gibco) with 15% fetal bovine serum (Gibco), MEM non-essential amino acids (Gibco), penicillin/streptomycin (Gibco), 550 µM 2-mercaptoethanol (Gibco), and 10 ng/ml of leukaemia inhibitory factor (LIF) (eBioscience).

### Antibody-bead coating

50 μl of Dynabeads G were washed twice in 200 μl of IP buffer (50 mM Tris-HCl (pH 7.0), 50 mM NaCl, 1% NP-40) and ∼10 μg of antibody were added. Antibodies used in this study: Spt6 (Cell Signalling – D6J9H); Ssrp1 (Biolegends – 10D1); TFIID (Santa Cruz – sc-273); Med14 (Invitrogen – PA5-44864), Total Pol II (anti-CTD, ab817 – Abcam), Sf1 (A303-214A; Bethyl).

### Nuclear extraction and DNase treament

A detailed protocol is available in the Supplementary Information (“NET-prism protocol”). Briefly, 10^8^ ES cells were used for each IP. It is important to split cells down to five batches of 2×10^7^ each when performing nuclei extraction. All extraction steps are performed on ice to avoid degradation of the nascent RNA. 2×10^7^ cells were treated with 200 μl of cytoplasmic lysis buffer (0.15% (vol/vol) NP-40, 10 mM Tris-HCl (pH 7.0), 150 mM NaCl, 25 µM α-amanitin (Epichem), 10 U RNasin Ribonuclease inhibitor (Promega) and 1× protease inhibitor mix (Thermo)) for 5 min on ice. Lysate was layered on 500 μl of sucrose buffer (10 mM Tris-HCl (pH 7.0), 150 mM NaCl, 25% (wt/vol) sucrose, 25 µM α-amanitin, 20 U RNasin Ribonuclease inhibitor and 1× protease inhibitor mix) and spun down for 5 min at 16,000g (4°C). The supernatant was carefully removed and nuclei were resuspended in 100 μl of DNase digestion buffer (1× DNase buffer (NEB), 25 µM α-amanitin, 20 U RNasin Ribonuclease inhibitor and 1× protease inhibitor mix) and further treated with 100 U of DNase I (NEB) for 20 min on ice. It is important for nuclei to be fully resuspended in the DNase digestion buffer. Non-resuspended nuclei are an indication of harsh cytoplasmic lysis conditions – In this case reduce the volume of cytoplasmic lysis buffer.

Chromatin-solubilised nuclei were spun down at 6,000g (4°C) for 2 min and the supernatant was carefully removed. Nuclei were further treated with 200 μl of nuclei lysis buffer (1% (vol/vol) NP-40, 20 mM HEPES (pH 7.5), 125 mM NaCl, 50 mM urea, 0.2 mM EDTA, 0.625 mM DTT, 25 µM α-amanitin, 20 U RNasin Ribonuclease inhibitor and 1× protease inhibitor mix) for 5 min on ice. Nuclei lysate was spun down at 18,500g (4°C) for 2 min and supernatants from five different batches were combined. Phosphatase inhibitor mix (x1) (Thermo) was implemented on all the above extraction steps for batches intended for Pol II S2ph and Pol II S5ph immunoprecipitations.

### Chromatin Immunoprecipitation (IP) and nascent RNA extraction

A detailed protocol is available in the Supplementary Information (“NET-prism protocol”). Briefly, combined supernatants from the previous step were incubated in a final 1/10 dilution in IP buffer for 2 hours at 4°C. For the sequential IP, a total Pol II antibody was used for 2 hours, followed by elution twice with 100 μl 2.5 mM CTD peptide (synthesized by Peptide Specialty Laboratories, Heidelberg, Germany; identical to Abcam ab17564) for 30 min. The eluate was further incubated with Ssrp1 antibody-coated beads for an additional 2 hours. Beads were washed 4 times with 1 ml of IP buffer and 700 μl of Qiazol (Qiagen) was directly added to the beads, followed by 140 μl of Chloroform. Samples were spun down and supernatant was ethanol precipitated (0.3M NaOAc, 2 μl Glycoblue). Concentration and size of nascent RNA was assessed by Nanodrop and TapeStation 2200, respectively. An IP from 10^8^ ES cells usually yields ∼200-1000 ng of nascent RNA. Assessment of RNA size is important in order to evaluate the fragmentation time during the library preparation.

### NET-prism library preparation

Two biological replicates were processed for each IP and library preparation. NET-prism libraries were prepared similarly to the human NET-seq protocol (Mayer and Churchman, 2016) with few modifications. The random barcode was ligated overnight at 16 °C to maximise ligation efficiency. Alkaline fragmentation of the ligated nascent RNA varies depending on the size of the RNA fragments obtained from each IP. IPs for Pol II S5ph, Pol II S2ph, Ssrp1, and Spt6 yielded large RNA fragments and therefore the ligated nascent RNA was fragmented until all RNA transcripts were within the range of ∼35-200 nucleotides. IPs for TFIID, and Mediator yielded fragments < 200 nt and therefore no fragmentation was performed. Maximum recovery of ligated RNA and cDNA was achieved from 15 % TBE-Urea (Invitrogen) and 10% TBE-Urea (Invitrogen), respectively, by adding RNA recovery buffer (Zymo Research, R1070-1-10) to the excised gel slices and further incubating at 70°C (1500 rpm) for 15 min. Gel slurry was transferred through a Zymo-Spin IV Column (Zymo Research, C1007-50) and further precipitated for subsequent library preparation steps. cDNA containing the 3’ end sequences of a subset of mature and heavily sequenced snRNAs, snoRNAs, and rRNAs, were specifically depleted using biotinylated DNA oligos (Mylonas et al.). Oligo-depleted circularised cDNA was amplified via PCR (9-12 cycles) and double stranded DNA was run on an 8% TBE gel. The final NET-seq library running at ∼150 bp was extracted and further purified using the ZymoClean Gel DNA recovery kit (Zymo Research). Sample purity and concentration was assessed in a 2200 TapeStation and further sequenced on a HiSeq 2500 Illumina Platform (Supplementary Table 2).

### NET-prism analysis

All the NET-prism fastq files were processed using custom Python scripts (https://github.com/BradnerLab/netseq) to align (mm10 genome) and remove PCR duplicates and reads arising from RT bias. Reads mapping exactly to the last nucleotide of each intron and exon (Splicing intermediates) were further removed from the analysis. The final NET-prism BAM files were converted to bigwig (1 bp bin), separated by strand, and normalized to ×1 sequencing depth using Deeptools (Ramírez et al., 2016) (v 2.4) with an “– Offset 1” in order to record the position of the 5’ end of the sequencing read which corresponds to the 3’ end of the nascent RNA. NET-seq/prism tags sharing the same or opposite orientation with the TSS were assigned as ‘sense’ and ‘anti-sense’ tags, respectively. Promoter-proximal regions were carefully selected for analysis to ensure that there is minimal contamination from transcription arising from other transcription units. Genes overlapping within a region of 2.5 kb upstream of the TSS were removed from the analysis. For the NET-seq/prism metaplots, genes underwent several rounds of k-means clustering in order to filter regions; in a 2kb window around the TSS, rows displaying very high Pol II occupancy within a <100 bp region were removed from the analysis as they represent non-annotated short non-coding RNAs. For Figure 1B, genes that displayed an RPKM > 1 for Total Pol II (n = 6,107) were used for metaplot profiling. Average Pol II occupancy profiles were visualised using R (v 3.3.0).

### Travelling ratio & Termination index

The travelling ratio is calculated via:

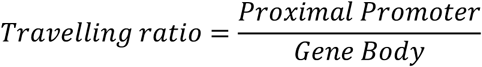

with Proximal Promoter defined as the Pol II coverage −30 bp and +250 bp around the TSS whereas Gene body region as the Pol II coverage +300 bp downstream of TSS and −200 bp upstream of TES.

The termination index is calculated via:

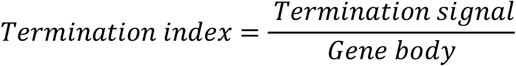

with the Termination signal defined as the Pol II coverage +2000 bp downstream of the TES whereas Gene body region as the Pol II coverage +300 bp downstream of TSS and −200 bp upstream of TES.

The reverse travelling ratio is calculated via:

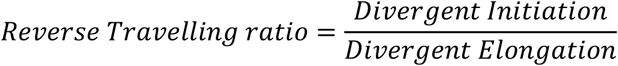

with Divergent Elongation being defined as the Pol II coverage between −1500 bp and −330 bp upstream of the TSS whereas Divergent Initiation being the Pol II coverage between −300 bp and −30 bp upstream of the TSS.

### ChIP-seq data processing

All ChIP-seq fastq files were aligned to the mm10 genome using Bowtie2 (v 2.2.6) with default parameters (Langmead and Salzberg, 2012). All BAM files were converted to bigwig (10 bp bin) and normalised to ×1 sequencing depth using Deeptools (v 2.4) (Ramírez et al., 2016). Duplicated reads were removed. Blacklisted mm9 co-ordinates were converted to mm10 using the LiftOver tool from UCSC and were further removed from the analysis. Average binding profiles were visualised using R (v 3.3.0).

### Mass spectrometry sample preparation

Independent ES cell cultures were grown in 10cm dishes. Per IP, 20×10^7^ cells were extracted, lysed, and nuclei were treated with DNase I as described above. The supernatant was incubated for 2 hours with a total Pol II antibody (ab817 – Abcam) or IgG (Cell Signalling) at 4°C. In total, four samples were prepared for each IP (Total Pol II, IgG). After thorough washing of beads with IP buffer, samples were incubated overnight at 37°C with Tris pH 8.8 and 300 ng Trypsin Gold (Promega). Peptides were desalted using StageTips (Rappsilber, Ishihama, & Mann, 2003) and dried. The peptides were resuspended in 0.1% formic acid and analysed using liquid chromatography - mass spectrometry (LC-MS/MS).

### LC-MS/MS analysis

Peptides were separated on a 25 cm, 75 μm internal diameter PicoFrit analytical column (New Objective) packed with 1.9 μm ReproSil-Pur 120 C18-AQ media (Dr. Maisch) using an EASY-nLC 1200 (Thermo Fisher Scientific). The column was maintained at 50°C. Buffer A and B were 0.1% formic acid in water and 0.1% formic acid in 80% acetonitrile. Peptides were separated on a segmented gradient from 6% to 31% buffer B for 45 min and from 31% to 50% buffer B for 5 min at 200 nl / min. Eluting peptides were analyzed on a QExactive HF mass spectrometer (Thermo Fisher Scientific). Peptide precursor m/z measurements were carried out at 60000 resolution in the 300 to 1800 m/z range. The top ten most intense precursors with charge state from 2 to 7 only were selected for HCD fragmentation using 25% normalized collision energy. The m/z values of the peptide fragments were measured at a resolution of 30000 using a minimum AGC target of 8e3 and 55 ms maximum injection time. Upon fragmentation, precursors were put on a dynamic exclusion list for 45 sec.

### Protein identification and quantification

The raw data were analyzed with MaxQuant version 1.6.0.13 (Cox and Mann, 2008) using the integrated Andromeda search engine (Cox et al., 2011). Peptide fragmentation spectra were searched against the canonical and isoform sequences of the mouse reference proteome (proteome ID UP000000589, downloaded December 2017 from UniProt). Methionine oxidation and protein N-terminal acetylation were set as variable modifications; cysteine carbamidomethylation was set as fixed modification. The digestion parameters were set to “specific” and “Trypsin/P,” The minimum number of peptides and razor peptides for protein identification was 1; the minimum number of unique peptides was 0. Protein identification was performed at a peptide spectrum matches and protein false discovery rate of 0.01. The “second peptide” option was on. Successful identifications were transferred between the different raw files using the “Match between runs” option. Label-free quantification (LFQ) (Cox et al., 2014) was performed using an LFQ minimum ratio count of 2. LFQ intensities were filtered for at least three valid values in at least one group and imputed from a normal distribution with a width of 0.3 and down shift of 1.8. The median value of the log2 LFQ intensities for the RNA Pol II IPs was used for the imputation of the missing values in the IgG IPs. Differential abundance analysis was performed using limma (Ritchie et al., 2015) (Supplementary Table 1).

### Enhancers and Super-enhancers

BED files containing typical enhancer and super-enhancer coordinates in mESCs were downloaded from Whyte et.al. (Whyte et al., 2013). Distal enhancers were defined as regions that are not overlapping with any annotated gene within a 2000 bp window. Only the distal enhancers that displayed an RPKM > 1 for Pol II were kept for subsequent analyses.

### Publicly available datasets

NET-seq (GSE90906) (Mylonas et al.), ChIP-seq (Pol II; GSE28247 (Handoko et al., 2011), Ssrp1; GSE90906 (Mylonas et al.), Spt6; GSE103180 (Wang et al., 2017), TFIID; GSE39237 (Ku et al., 2012), H3K27Ac (Encode Consortium – E14 cell line))

## DATA AVAILABILITY

Data have been deposited in Gene Expression Omnibus (GEO) under accession numbers GSE 107257.

## ACKNOWLEDGEMENT

We are grateful to Ilian Attanassov of the Max Planck Institute for Biology of Ageing Proteomics Core Facility for Mass Spectrometry Analysis. Sequencing was performed at the Max Planck Genome core centre in Cologne and data analysis was done on servers of the GWDG, Göttingen and the MPI-AGE cluster. We thank members of the Tessarz laboratory for discussion and comments on the manuscript.

## SUPPLEMENTARY DATA

Supplementary Data are available at online.

## SUPPLEMENTARY FIGURES

**Supplementary Figure 1:**
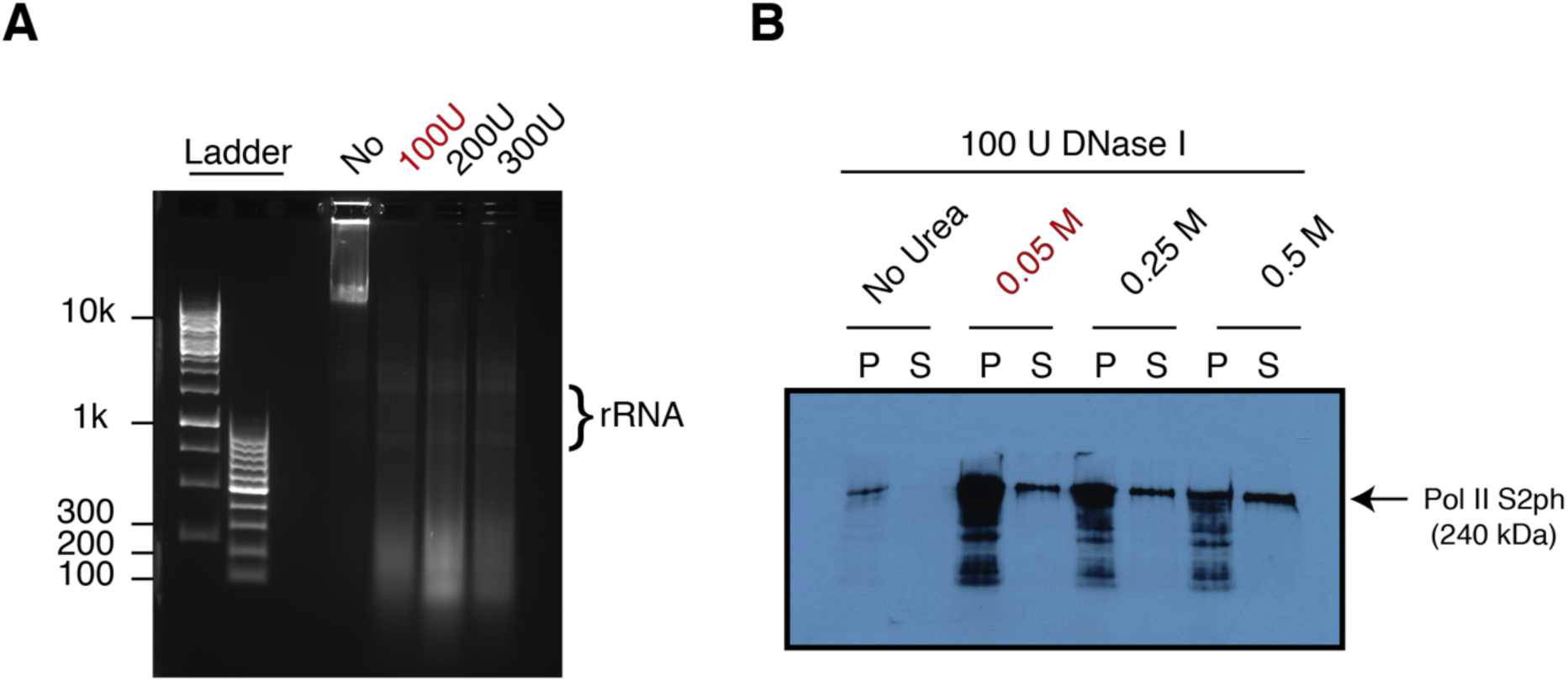
Release of Pol II from chromatin via DNase I treatment. (A) Agarose gel (1%) showing the effect of different concentrations of DNAse I on DNA fragmentation. No difference is observed among treated conditions. (B) Western blot assessing Pol II release after treatment of 2×10^7^ mouse ES cells with different Urea concentrations (0, 0.05 M, 0.25 M, 0.5 M) in the presence of 100U of DNase I. The conditions highlighted in red (0.05 M Urea & 100U DNAse I) were used for the generation of all NET-prism libraries (P; Pellet, S; Supernatant).

**Supplementary Figure 2:**
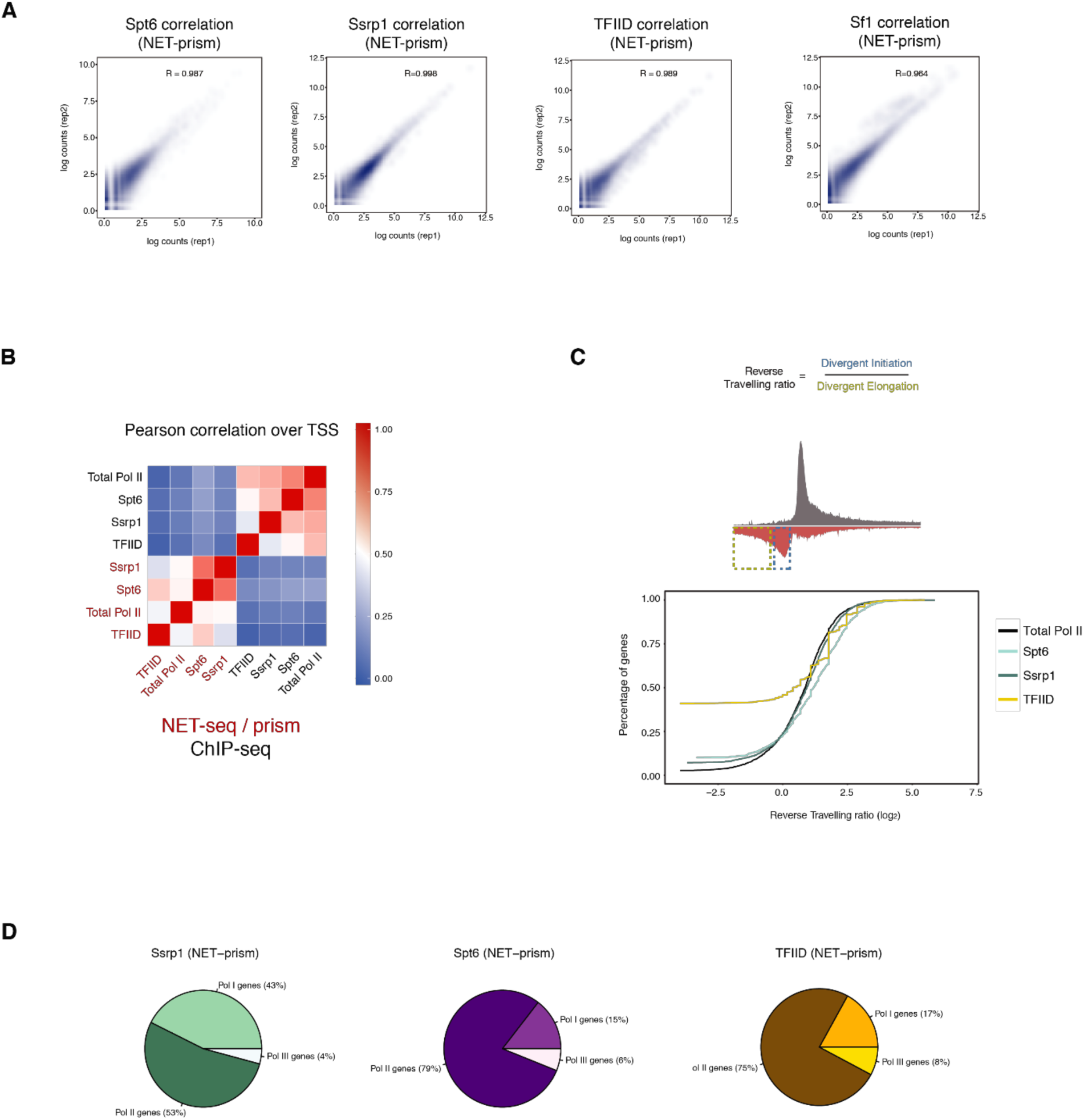
Transcriptional activity assessed by NET-prism. (A) Correlation plots assessing reproducibility between replicates of Spt6, Ssrp1, TFIID, and Sf1 libraries (NET-prism). R corresponds to Pearson correlation. (B) Correlation heatmap of all NET-seq/prism (red) and ChIP-seq (black) libraries over promoter regions (± 500 bp around TSS) of all uniquely annotated genes (n = 12,737). (C) Top: Schematic diagram for the calculation of the reverse travelling ratio. Bottom: Measurement of the Reverse Travelling ratio over uniquely annotated bidirectional promoters (n = 2,254) for the indicated NET-prism libraries. (D) Distribution of reads mapping to Pol I, Pol II, Pol III transcribed RNAs for Ssrp1, Spt6, and TFIID NET-prism libraries in mESCs.

**Supplementary Figure 3:**
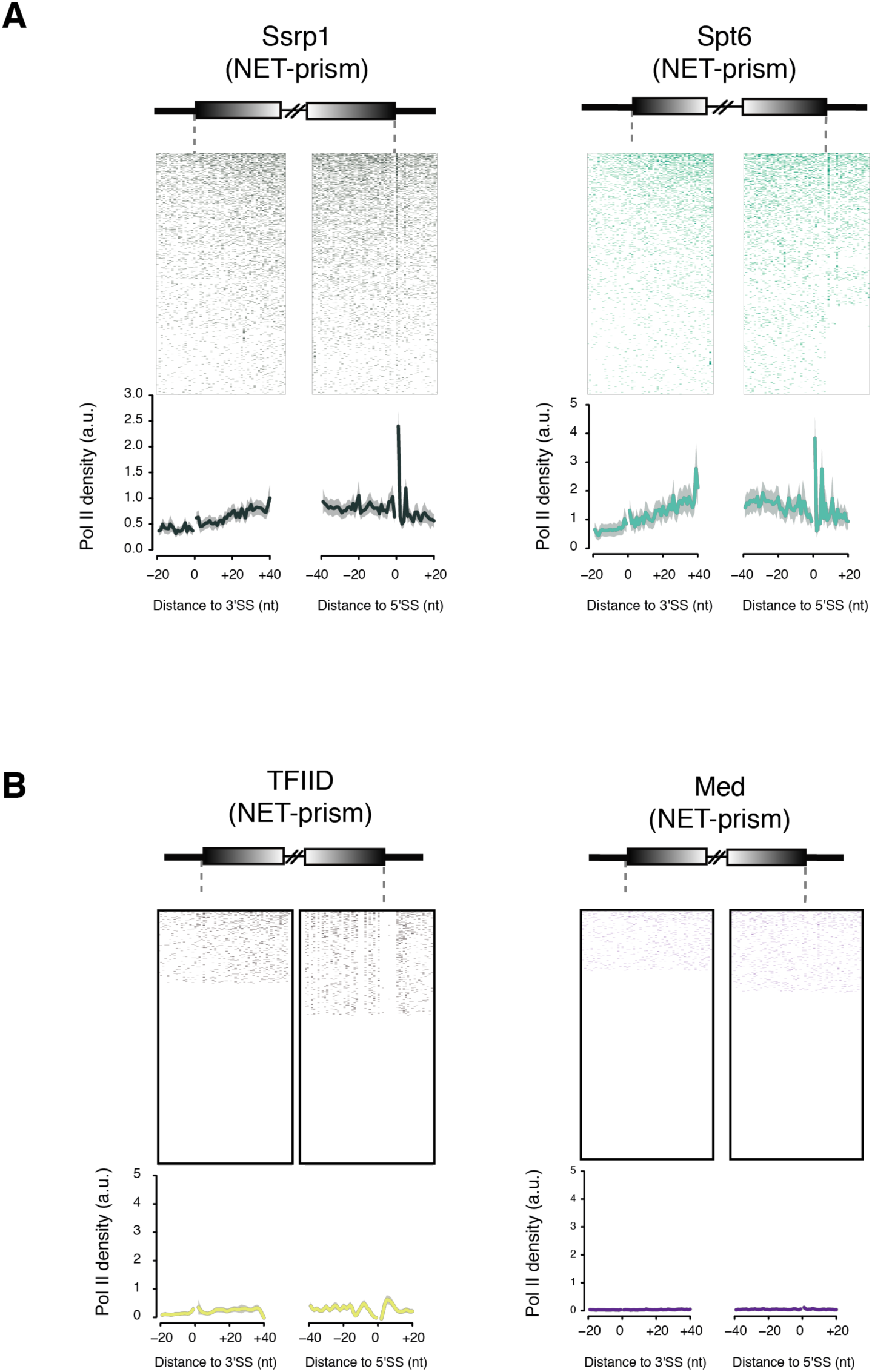
No association of the PIC complex with co-transcriptional splicing. (A) Heatmaps and metaplots assessing polymerase pausing for Ssrp1, and Spt6 over exon boundaries (n = 2,586). Solid lines indicate the mean values, whereas the shading represents the 95% confidence interval. (B) Heatmaps and metaplots assessing polymerase pausing for TFIID, and Mediator over exon boundaries (n = 2,586). Solid lines indicate the mean values, whereas the shading represents the 95% confidence interval.

**Supplementary Figure 4:**
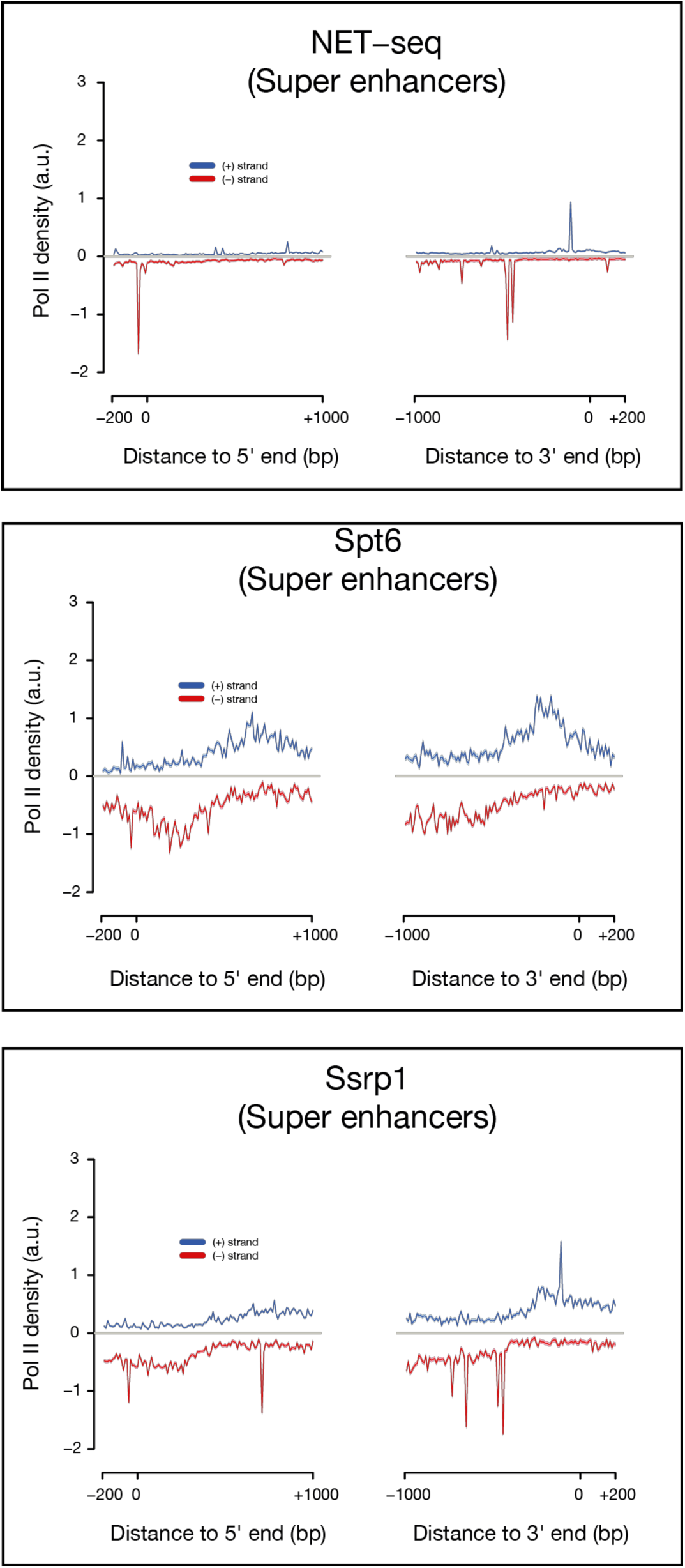
Unique patterns of transcriptional pausing over super-enhancers. Metaplots assessing polymerase pausing for total Pol II, Pol II S5ph, Ssrp1, and, Spt6 over super-enhancers (n = 226). Solid lines indicate the mean values, whereas the shading represents the 95% confidence interval.

## NET-prism PROTOCOL

### Quantitative purification of RNA polymerase by cell fractionation • TIMING 45 min

Cell fractionation is performed on ice or at 4 °C, with buffers freshly prepared on the same day. All buffers are precooled on ice before use.

Use RNase-free reagents and equipment.

**<u>Preparation of antibody-conjugated beads</u>** • TIMING 2 h (or overnight)

1| Wash 50 µl of Dynabeads protein G with 1 ml of ice-cold IP buffer by vortexing at room temperature: repeat once more.

3| Resuspend the Dynabeads in 50 µl of ice-cold IP buffer.

4| Add ∼10 µg of the appropriate antibody to washed beads. IgG can be used as control (optional).

5| Incubate the mixture on a tube rotator (12 r.p.m.) in a cold room for at least 2 h.

7| Briefly spin down tubes and place on ice until use.

**1|** Grow cells in a 15 × 2.5 cm dish in DMEM containing 10% (vol/vol) FBS, 100 U/ml penicillin and 100 µg/ml streptomycin until they are 90% confluent

**2|** Wash the cells twice with 10 ml of PBS buffer.

**CRITICAL STEP** Unless otherwise stated, perform all washing steps on ice.

**3|** Scrape or trypsinise cells into 1 ml of PBS buffer.

**4|** Determine the cell number by counting using a cell counter. Use 2×10^7^ cells as input for each cell fractionation. You will need in total 80-100×10^7^ cells for each IP.

**5|** Wash each sample of 2× 10^7^ cells with 10 ml of PBS.

**6|** Collect cells by centrifugation at 500g for 2 min at 4 °C.

**7|** Remove the supernatant by aspiration.

**CRITICAL STEP** It is important to remove the supernatant completely at this step. If the supernatant is not completely removed, the cytoplasmic lysis buffer (Step 8) will be diluted, which affects the cell lysis efficiency.

**8|** Add 200 µl of cytoplasmic lysis buffer and transfer it to an RNase-free 1.5-ml microcentrifuge tube. Cut a 1,000-µl pipette tip, and use it to pipette the sample up and down ten times.

**9|** Incubate the cell lysate on ice for 5 min.

**10|** Using a cut 1,000-µl pipette tip, layer the cell lysate onto 500 µl of sucrose buffer.

**11|** Collect cell nuclei by centrifugation at 16,000g for 10 min at 4 °C.

**12|** Remove the supernatant.

**13|** Add 100U of DNAse I in 100 µl of DNAse buffer (NEB) and place on ice for 20 min. Stop reaction by adding EDTA to a final concentration of 5 mM.

**14|** Wash nuclei with 800 µl of nuclei wash buffer.

**15|** Collect washed nuclei by centrifugation at 7,000g for 1 min at 4 °C.

**16|** Add 200 µl of nuclei lysis buffer, mix by pulsed vortexing and incubate the mixture on ice for 5 min.

**17|** Centrifuge the mixture at 18,500g for 2 min at 4 °C.

**18|** Combine supernatants from different fractionations (Final volume: 800 µl or 1000 µl)

**19|** Transfer to a 15ml falcon tube and add IP buffer (7.2ml or 9ml) to a final 1/10 dilution. Add RNase/protein inhibitors.

**20|** Add antibody–conjugated beads and mix immediately.

**21|** Incubate the samples on a tube rotator (12 r.p.m.) for 2 h in the cold room.

**22|** Spin the samples at 300g for 5 min at 4 °C.

**23|** Aspirate supernatant. Leave only 1ml behind and transfer to a new 1.5ml tube.

**24|** Place the tube on a magnetic rack and leave it for 1 min. Remove the remaining supernatant.

**27|** Wash the beads with 1 ml of ice-cold IP buffer by inverting tube, and repeat Step 24 three times (total 4 times) in the cold room, taking care to remove the supernatant after the final wash.

**28**| Add 700 µl of TRIzol reagent directly to the beads and mix well by pipetting.

**29**| Add 150 µl of chloroform. Mix well and leave for 2 min at room temperature.

**30**| Centrifuge at 18,000g for 30 min at 4 °C.

**31**| Carefully transfer the upper transparent layer (∼300 µl) to a new tube without disrupting the phenol layer.

**32**| Add 900 µl 100% ethanol, 30 µl NaOAc (3M), and 2 µl GycoBlue.

**33**| Incubate at −80 °C for 1h.

**34**| Centrifuge at 18,000g for 30 min at 4 °C.

**35**| Remove supernatant and wash pellet once with 1ml 100% ethanol.

**36**| Centrifuge at 18,000g for 2 min at 4 °C.

**37**| Remove supernatant and wash pellet once with 1ml 70% ethanol.

**38**| Centrifuge at 18,000g for 2 min at 4 °C.

**39**| Completely remove supernatant and let pellet air dry on ice for 5 min.

**40**| Resuspend pellet in 13 µl of RNAse-free water.

**41**| Run 1 µl on a Tapestation to assess RNA size and concentration. Typical concentration is between 500 – 1000 ng/ µl. IPs for elongations factors (Spt6) yield large RNA fragments (>2000 nt) whereas PIC components yield lower (< 200 nt).

**42**| Parse samples into the same NET-seq library as previously described (Mayer & Churchman, 2016: *Nature Protocols*). Make the appropriate modifications as described below to allow a more efficient adaptor ligation and material recovery after ligation and cDNA synthesis.

- The random barcode was ligated overnight at 16 °C to maximise ligation efficiency.
- Alkaline fragmentation of the ligated nascent RNA varies depending on the size of the RNA fragments obtained from each IP. IPs for Pol II S5ph, Pol II S2ph, Ssrp1, Spt6, and Sf1 yielded large RNA fragments and therefore the ligated nascent RNA was fragmented until all RNA transcripts were within the range of ∼35-200 nucleotides. IPs for TFIID, and Mediator yielded fragments < 200 nt and therefore no fragmentation was performed.
- Maximum recovery of ligated RNA and cDNA was achieved from 15 % TBE-Urea (Invitrogen) and 10% TBE-Urea (Invitrogen), respectively, by adding RNA recovery buffer (Zymo Research, R1070-1-10) to the excised gel slices and further incubating at 70°C (1500 rpm) for 15 min. Gel slurry was transferred through a Zymo-Spin IV Column (Zymo Research, C1007-50) and further precipitated for subsequent library preparation steps.

### Buffers (×1 reaction)

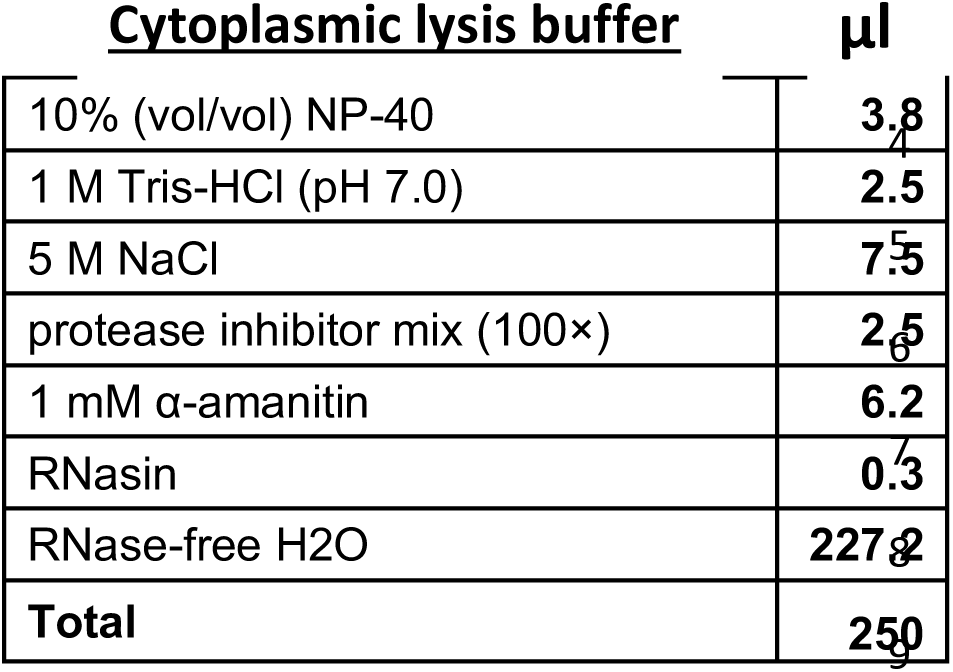

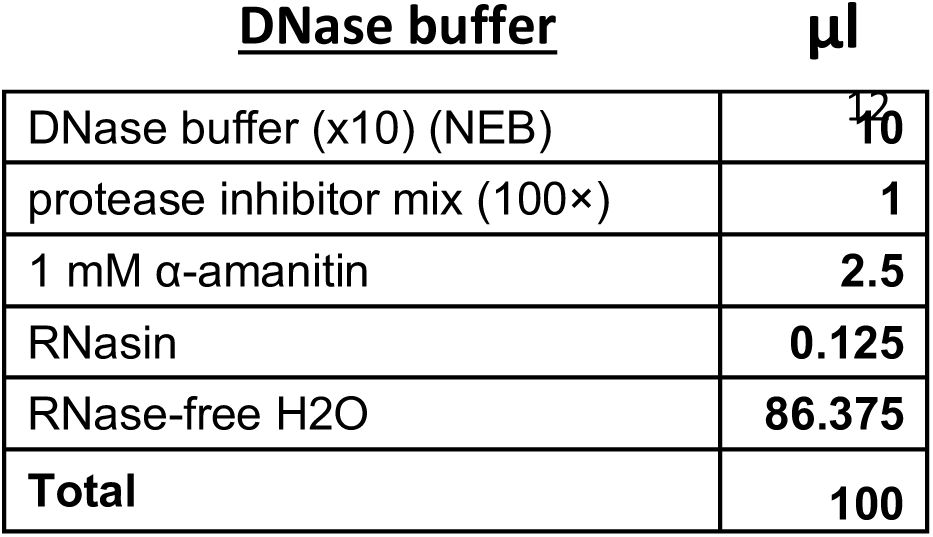

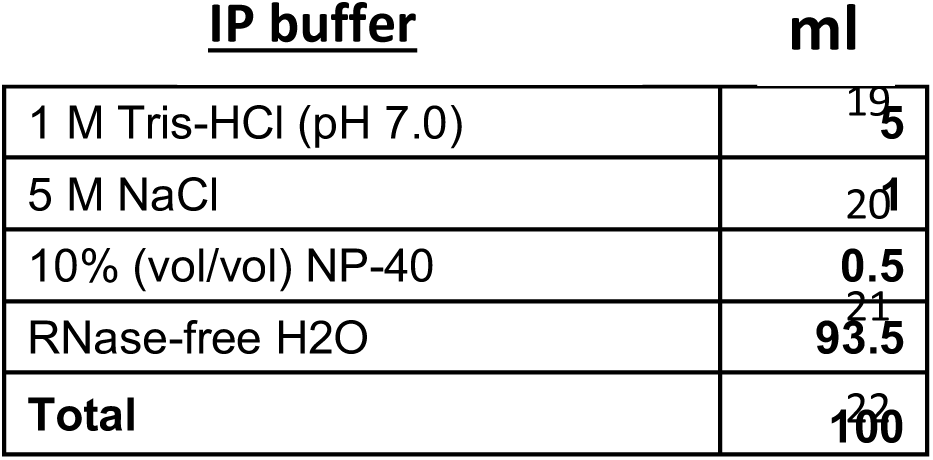

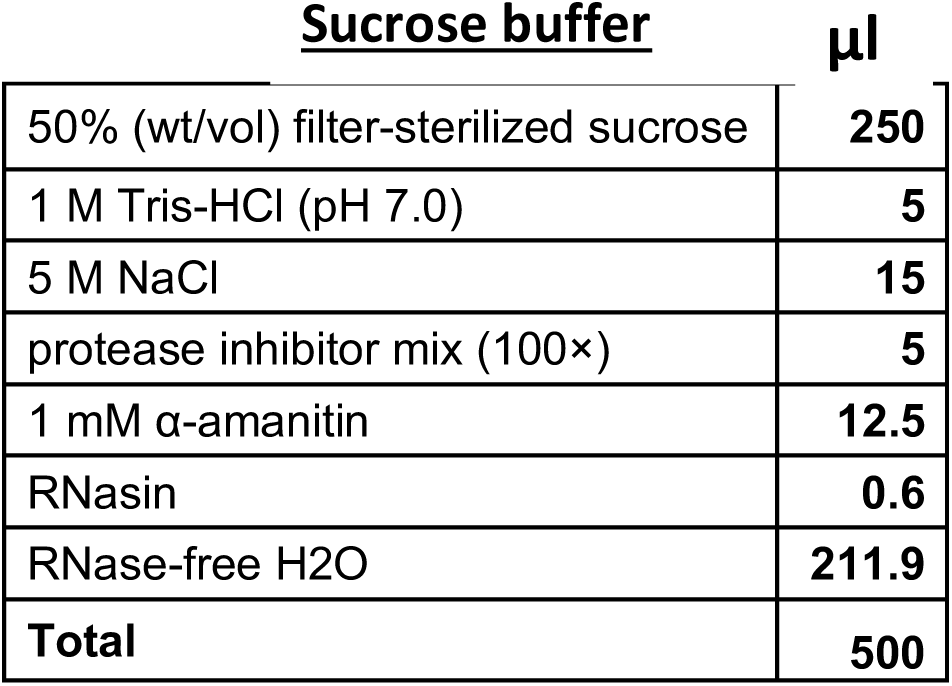

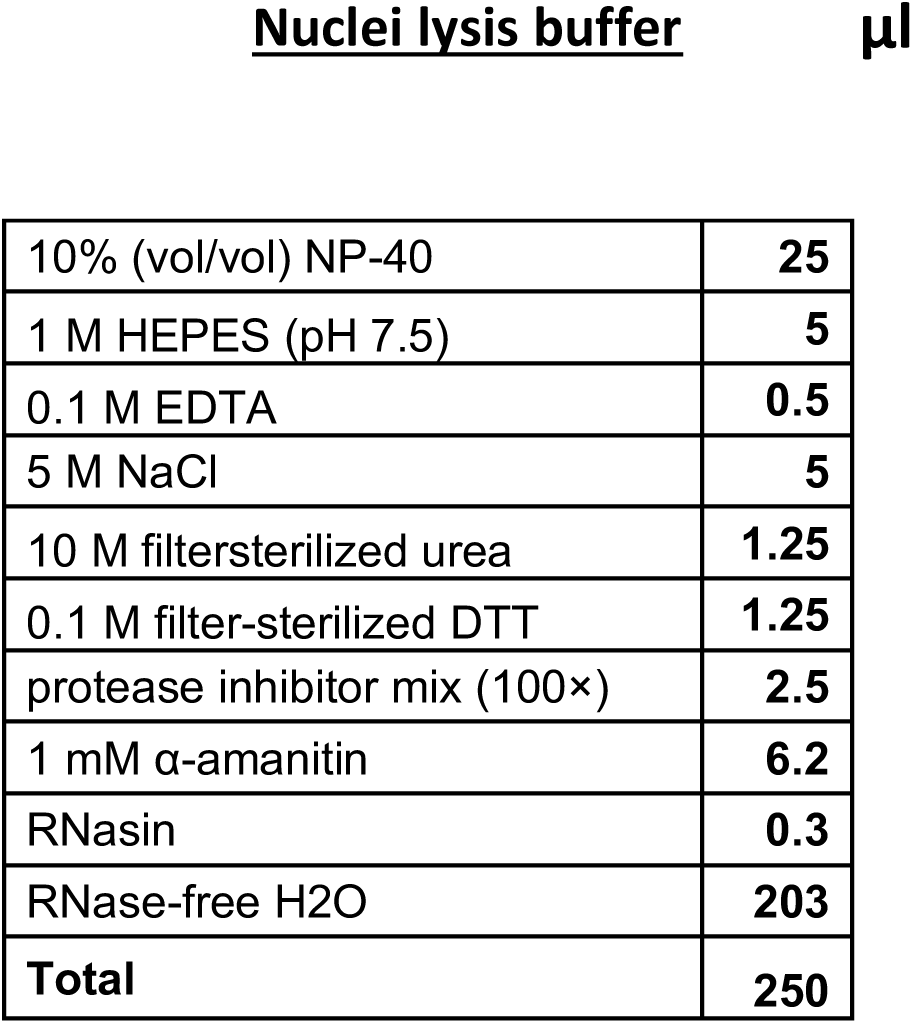

